# First data-driven approach to using individual cattle weights to estimate mean adult dairy cattle weight in the UK

**DOI:** 10.1101/270702

**Authors:** Hannah E. Schubert, Sarah Wood, Kristen K. Reyher, Harriet L. Mills

**Affiliations:** Bristol Veterinary School, University of Bristol, Langford House, Langford, Bristol, BS40 5DU, UK.; MRC Integrative Epidemiology Unit, Bristol Medical School, University of Bristol, Oakfield House, Oakfield Road, Bristol, BS8 2BN, UK

**Keywords:** Dairy cattle, weight, automatic milking systems, antimicrobial usage, medicine dosing

## Abstract

**Introduction:** Knowledge of accurate weights of cattle is crucial for effective dosing of individual animals with medicine and for reporting antimicrobial usage metrics, amongst other uses. The most common weight for dairy cattle presented in current literature is 600 kg, but this is not evidenced by data. For the first time, we provide an evidence-based estimate of the average weight of UK dairy cattle to better inform decisions by farmers, veterinarians and the scientific community.

**Methods:** We collected data for 2,747 dairy cattle from 20 farms in the UK, 19 using Lely Automatic Milking Systems with weigh floors and 1 using a crush with weigh scales. These data covered farms with different breed types, including Holstein, Friesian, Holstein-Friesian and Jersey, as well as farms with dual purpose breeds and cross-breeds. Data were used to calculate a mean weight for dairy cattle by breed, and a UK-specific mean weight was generated by scaling to UK-specific breed proportions. Trends in weight by lactation number, DIM and production level were also explored using individual cattle-level data.

**Results:** Mean weight for adult dairy cattle included in this study was 617 kg (standard deviation (sd) 85.6 kg). Mean weight varied across breeds, with a range of 466 kg (sd=56.0 kg, Jersey) to 636 kg (sd=84.1, Holsteins). When scaled to UK breed proportions, the estimated mean UK dairy cattle weight was 620 kg. Overall, first-lactation heifers weighed 9% less than cows. Mean weight declined for the first 30 days post-calving, before steadily increasing. For cattle at peak production, mean weight increased with production level.

**Conclusions:** This study is the first to calculate a mean weight of adult dairy cattle in the UK based on on-farm data. Overall mean weight was higher than that most often proposed in the literature (600 kg). Evidence-informed weights are crucial as the UK works to better monitor and report metrics to monitor antimicrobial use and are useful to farmers and veterinarians to inform dosing decisions.

## INTRODUCTION

Average weights of dairy cattle in the UK are not well defined. Scientific papers, reports and guidelines present a wide range of adult dairy cattle weights. A literature review demonstrated a range from 425 kg (EU estimated “average weight at time of treatment” (European Surveillance of Veterinary Antimicrobial Consumption)) to 680 kg (USA) (Pol and Ruegg, 2007) (Table 1). Additionally, the weights used in current literature are commonly either “estimated”, without clear evidence, or cited from another source (usually equally lacking in evidence). Average cattle weight would also be expected to vary with breed (DairyCo, 2005) and between populations (Collineau et al., 2017) (e.g. countries, due to different compositions of herds nationally), but this is rarely accounted for in the literature.

**Table 1.** Cattle weights as presented in recent literature. This list is not the result of a systematic literature search but does reflect the most commonly used and cited weights. Note that the majority of these have been defined for measuring antimicrobial usage

| Reference | Weights (kg) |  |  |  |  |  | Comments |
| --- | --- | --- | --- | --- | --- | --- | --- |
|  | Milking<br>cow | Dairy<br>heifers<br>>24<br>months | Dairy<br>heifers<br>12-24<br>months | Dairy<br>heifers<br>weaning<br>- 12<br>months | Pre<br>weaned<br>calves | Other |  |
| (Montforts, 2006) | 425 |  |  | 200 | 200 |  | Not justified with a reference or data. |
| Methodology for<br>Determining Antibiotic<br>Use (European<br>Surveillance of<br>Veterinary<br>Antimicrobial<br>Consumption)<br>(ESVAC) | 425 | 425 | 200 | 140 | 140 | Bullocks<br>and<br>bulls:<br>425 | Average weight at time of treatment, based<br>on the assumption that younger animals are<br>more likely to have antimicrobial treatment.<br>Weights derived from a committee of<br>European experts, citing (Montforts, 2006).<br>Cited widely in other literature. |
| UK-VARRS<br>(Veterinary Medicines<br>Directorate & Animal<br>and Plant Health<br>Agency, 2015) | 425 |  |  | 140 |  |  | Average weight at time of treatment. Cites<br>(European Surveillance of Veterinary<br>Antimicrobial Consumption) and (Montforts<br>et al., 1999) (note, this Montforts paper<br>actually gives a weight of 600 kg, we assume<br>the correct citation is (Montforts, 2006)). |

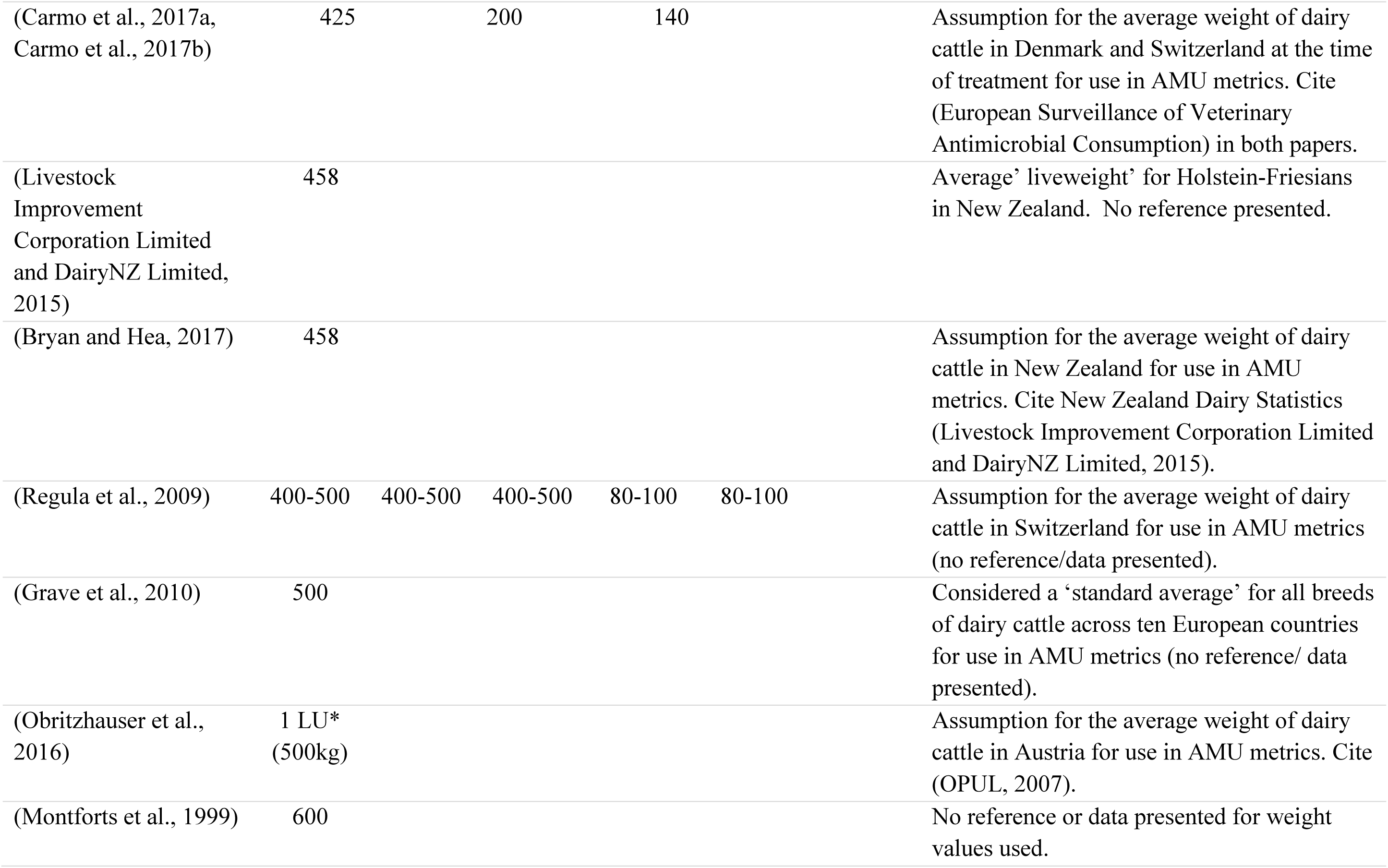

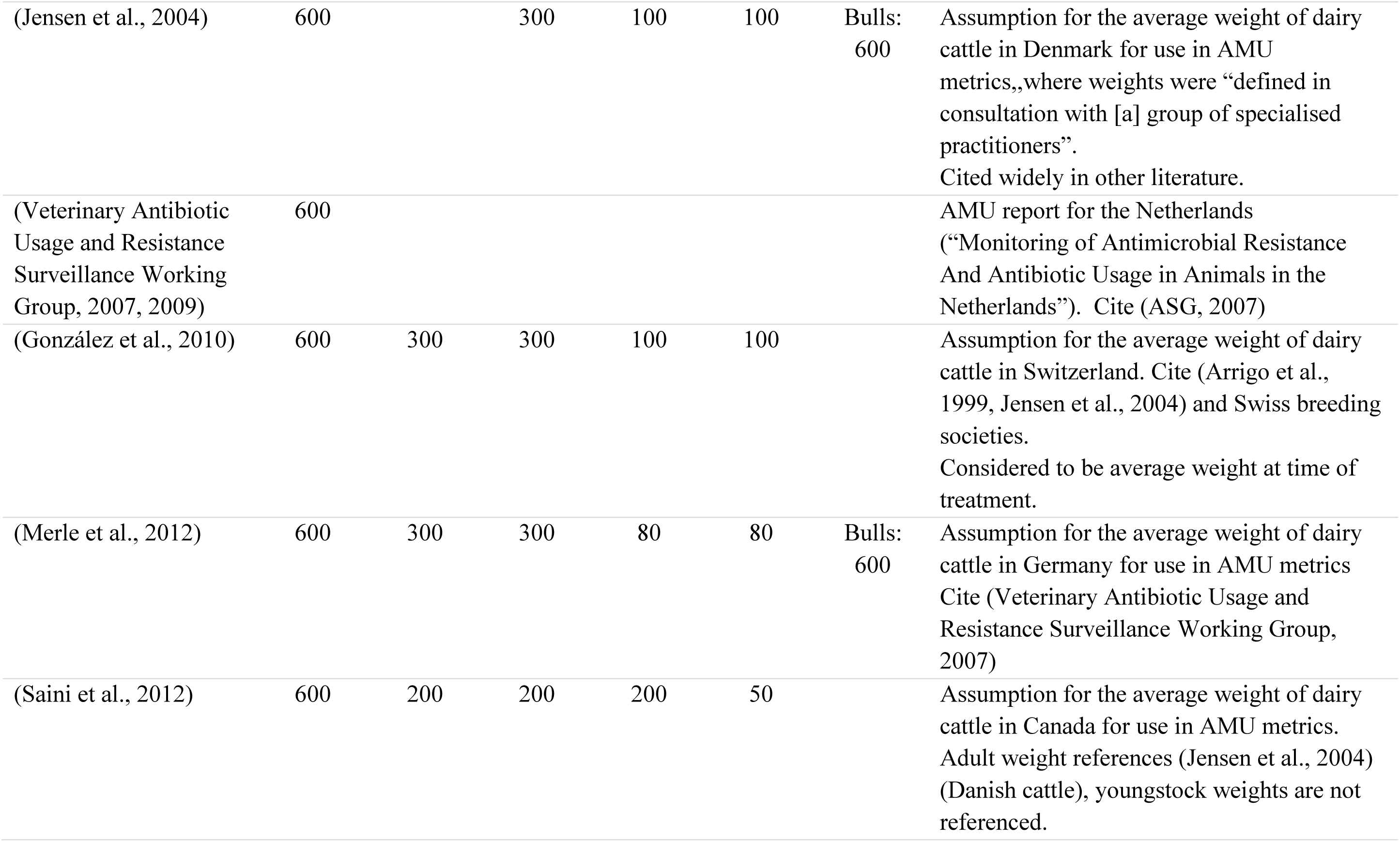

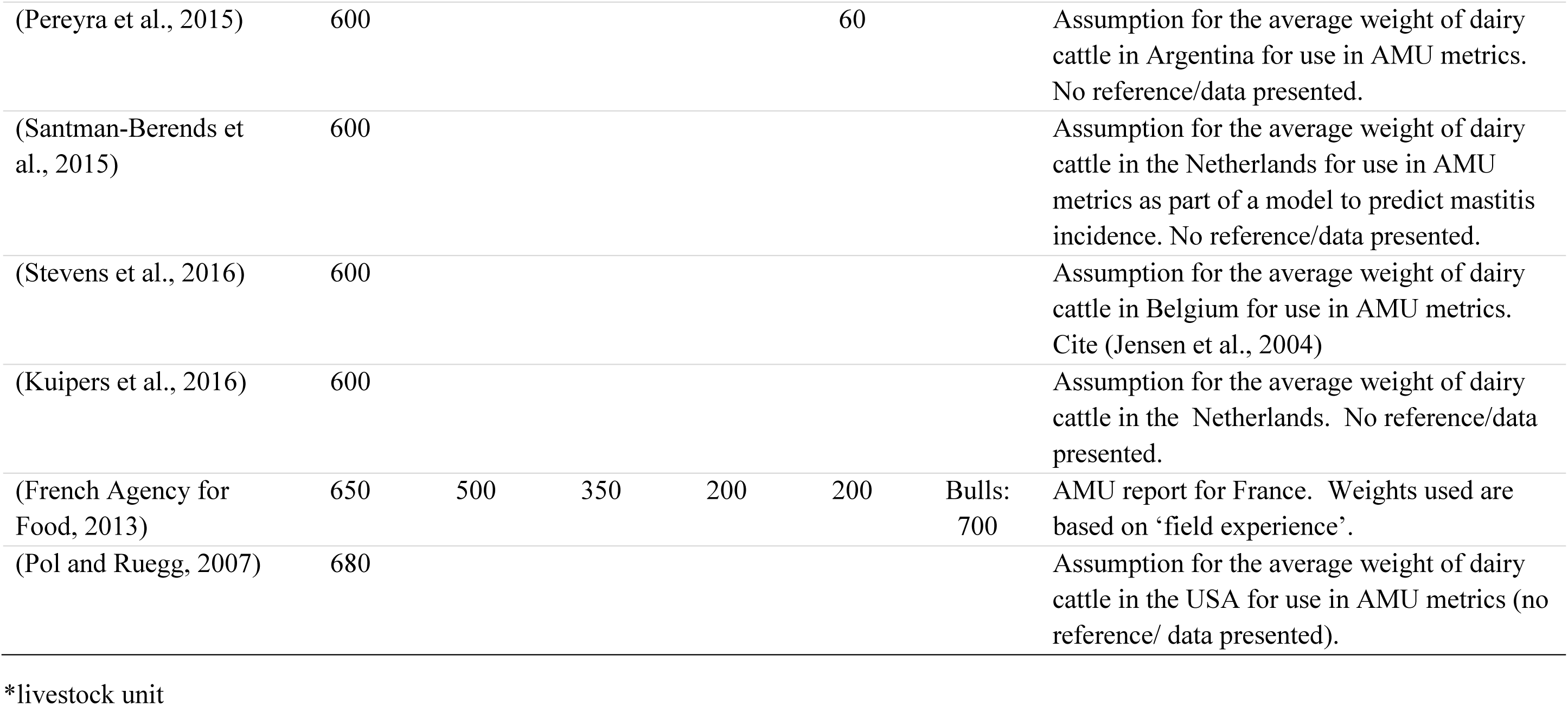

Many medicine doses should be calibrated to the estimated weight of the cattle being treated. Using incorrect weights may lead to incorrect dosing, which could prove ineffective or potentially dangerous. This is particularly true of antimicrobials where an underdose could fail to completely clear the infection, a problem which has been linked to the risk of resistance developing (Roberts et al., 2008). Additionally, metrics for reporting antimicrobial use (AMU, for example mg/kg or daily dose metrics, (Mills et al., 2017)) commonly require the total weight of the animals at risk of treatment to be included in the calculation, giving a measure which accounts for the total kg. If the included weight is too high or too low this could lead to the metric under- or over-representing the actual use of the antimicrobials and confound comparison across farms or countries.

For the purpose of treating cattle with the appropriate dose of medicine, visually estimated weight is usually relied upon. However, it has previously been shown that visual estimates of cattle weights vary in accuracy compared with estimates from heart girth tape measurements, with under- and overestimation at the extremes of the weight scale (Wood et al., 2015). Visual estimates may also be influenced by expectations of weight, which can also vary widely. For example, we asked 15 farm vets in practices across South West England to estimate the average weight of a UK ‘Holstein-Friesian milking cow’, resulting in a range of 525-775 kg and a mean of 678 kg.

Additionally, weight estimates based on body measurements of cattle (e.g. Schaeffer’s formula (Sastry et al., 1983)) or use of weigh tapes (Heinrichs et al., 2007) have been shown to deviate from true weights (Wangchuk et al., 2017). More accurate measures can be obtained from scales such as weigh crushes or weigh floors.

Some Automatic Milking Systems (AMS) have a weigh floor that records cattle weight at every milking (e.g. Lely, https://www.lely.com/gb/). These are predominantly used to monitor changes in weight and draw the stockperson’s attention to abnormal losses or gains (for example, Lely suggest a daily weight loss of 0.8% would require attention (**Lely**)). These weigh floors have been used in previous studies to monitor cattle weight change over time (van der Tol and van der Kamp, 2010, Podlahová et al., 2011). They are precisely calibrated at installation, and are cleaned and set to zero at every service (approximately 7 times over every 2-year period). Equipment is also widely available for weighing cattle through a handling crush.

In order to determine mean UK adult dairy cattle weights for use by farmers, veterinarians and the scientific community, we used data collected from 20 UK farms (19 from farms using Lely AMS and 1 farm using a crush with weigh scales). We also used these data to establish mean breed weights and to explore trends in weight by lactation number, DIM and overall milk production.

## MATERIALS AND METHODS

### Data collection

We collected data from 20 UK farms: 19 of these farms used Lely AMS and were recruited through Lely - 10 from Cornwall and Devon, 6 from Somerset and 3 from different areas of the UK. Lely emailed the farms from Devon, Cornwall and Somerset asking farmers to give permission to Lely to access the farm’s AMS data for a single day (See Appendix 1). Data from the other 3 farms came from another study to which Lely had contributed. Farmers were asked to calibrate the AMS weigh floor scales (“calibrate” being the term used by Lely to describe the following: clean scales and remove any trapped stones, then select “tare scale” on the control screen) and contact Lely to let them know this had been done (by text message). Lely then remotely downloaded a report from the farm’s AMS. The 20th farm was recruited directly and cattle were weighed using a crush with a weigh bar and digital scales. This farm in Devon with a Jersey herd was included, despite the different weighing method, for maximum representation across breeds. All cattle from the milking herd were weighed. An operator whose weight was known stood on the scales prior to use to check for accuracy, and the scales were set to zero between cattle if necessary.

Datasets from Lely were fully anonymised before we received them and contained the following relevant variables: *Animal Number*, *Robot*, *Date Time*, *Lactation No*., *Lactation days*, *Weight at Calving*, *Weight*, *Weight Avg*., *Weight Avg*. *Dev*. *and Day Production*. The variable “Weight” was the weight at last milking and “Weight Avg.” was the mean weight for that animal over the last 3 milkings (from the current lactation). “Weight at Calving” was the very first weight recorded after calving for the current lactation. “Weight Avg.” was used for all calculations. The dataset acquired using a crush was anonymised at animal level and contained the following variables: *Animal Number*, *Date*, *Lactation No*., *Lactation days and Weight*.

Breeds were assigned at the farm level by the farmer (Holstein = 7 farms, Friesian = 2 farms, Holstein-Friesian = 8 farms, Jersey = 1 farm, Cross-breed = 1 farm, Other breed = 1 farm; Table S1). All farms were all-year round calving which meant a full range of lactation stages were included.

### Data cleaning

Farm datasets from Lely contained data for all milking cattle registered to that farm at the time the report was taken. This included the last weight and production measurements for cattle that had not been milked recently. Cattle not weighed recently were likely to be dry, therefore the measurement was likely to be from the end of their previous lactation; including these would have caused an over-representation of late lactation cattle. Additionally, extreme dates may have indicated that the electronic collars used by the AMS for identification may have been broken, or that the system was not updated to indicate that an animal was removed from the herd. Therefore, for each farm, we used only data from the date with the most cattle milked/measured and the immediate week preceding (Table S1). Entries with missing weight or missing date were also removed; only 1 entry per animal was kept.

At the Jersey farm, data was excluded if the scales were not set to zero in between cattle.

### Representativity of data

To check that the cattle used in this study were representative of the UK herd, we obtained data on the proportion of heifers, mean lactation number and mean herd size. These data came from all UK herds that milk record with National Milk Records (NMR). The proportion of heifers in the NMR data was compared to our sample using a chi-square test for equal proportions. As the herds included in our dataset will be included in the herds provided by NMR, only simple comparisons were possible for mean lactation number and mean dairy herd size (a t-test would have required distinct subsets).

### Data analysis

We calculated the distribution and descriptive statistics for mean weights of cattle for the following breed categories: Holstein, Friesian, Holstein-Friesian, Cross-breed, Jersey, Other breed. Weights were calculated overall (for all cattle) and split into first lactation only (heifers) and second lactation onwards (cows). Overall mean weights and heifer and cow weights were compared across breeds using t-tests. Mean weights of heifers and cows for each breed and for the dataset as a whole were also compared using t-tests.

Additionally, the mean weight for cattle in each day of lactation (overall, and split into heifers and cows) was calculated and plotted to identify any trends over lactation. The correlation between mean weight and daily milk production was calculated. As milk production is known to vary across lactation, this analysis was repeated with only cattle considered to be in peak production (20-60 days into lactation).

Data analysis and graphics were generated using the statistical computing package R (https://www.r-project.org/).

### Estimated average weight for the UK

By comparing the proportion of each breed within our dataset to the proportion in the UK population (using data provided by the British Cattle Movement Service (BCMS), Table S3), we calculated an estimated average adult dairy cattle weight for the UK. Breeds reported by BCMS were grouped into categories (Table S3 & S4) aligned with the breeds for our data. To estimate a UK national average weight, mean weights by breed category calculated from our data were scaled according to the representation of that category within BCMS data.

### Calibration checks

For each farm using Lely AMS, we calculated the distribution of weights for each of the farm’s individual AMS units and checked for any which showed unexpected deviation from the overall mean for that farm and breed using t-tests. Additionally, we collected 6 days of weight data directly from the farmer from the largest farm (with the highest number of AMS units) over a 1-week period. These data were used to check the calibration accuracy of the individual weigh floors by comparing the mean and distribution of weights each day using t-tests.

## RESULTS

### Data description

The original datasets included 3,106 cattle; after cleaning, 2,747 cattle remained (i.e. 11.5% of cattle were excluded due to dates outside of range, missing date or weight information or our inability to obtain an accurate weight). Table 2 presents summary statistics for the cattle included in the study. Just under a third of cattle were in their first lactation. On the date of sampling, mean production was 33 L (Table 2).

**Table 2.** Summary data for 20 farms and 2,747 cattle remaining after cleaning for date and missing data was performed

| **Breakdown of cattle** |  | **N (%)** |
| --- | --- | --- |
| Total farms |  | 20 |
| By breed* | Holstein | 7 (35.0%) |
|  | Friesian | 2 (10.0%) |
|  | Holstein-Friesian | 8 (40.0%) |
|  | Jersey | 1 (5.0%) |
|  | Cross-breed | 1 (5.0%) |
|  | Other | 1 (5.0%) |
| Total cattle |  | 2,747 |
| By breed* | Holstein | 1,099 (40.0%) |
|  | Friesian | 130 (4.7%) |
|  | Holstein-Friesian | 1,099 (40.0%) |
|  | Jersey | 170 (6.2%) |
|  | Cross-breed | 197 (7.2%) |
|  | Other | 52 (1.9%) |
| By lactation number | 1 (heifers) | 857 (31.2%) |
|  | 2+ (cows) | 1,890 (68.8%) |
| **Summary statistics of key properties** |  | **Mean (SD)** |
| Number of cattle per farm** |  | 137.3 (sd=74.9) |
| Lactation number |  | 2.7 (1.8) |
| DIM |  | 174.3 (116.2) |
| Production data (litres)*** |  | 32.7 (11.2) |

\*Note that breed is assigned at farm level.
\*\*Note this is a mean across farms after some cattle were removed due to cleaning, actual mean herd size was 155 (cows currently in milk only)
\*\*\*Production data was not available for the Jersey cattle

### Representativity of data

Data provided by NMR on all dairy cattle in UK herds indicated that the mean proportion of heifers within a herd nationally was 29.1% (95% CI [29.0%, 29.2%]), compared to 31.2% (95% CI [29.5 %, 33.0 %]) within our dataset (Table 2). The mean lactation number within herds nationally was 2.8, compared to 2.7 within our dataset (Table 2). The mean number of cows in milk nationally was 155, compared to 155 within our dataset (Table 2).

### Data analysis

The cattle within this dataset had an overall mean weight of 617.3 kg (standard deviation 85.6 kg, median 620 kg) across all breeds and including both heifers and cows (Table 3). Heifers were on average 9.0% lighter than cows (Figure 1A) with mean weight 578.0 kg for heifers and 635.2 kg for cows (t-test: p<0.05). Jersey cattle were 25.8% lighter than the overall mean weight for all other breeds (465.7 kg compared to 627.3 kg).

**Figure 1:**
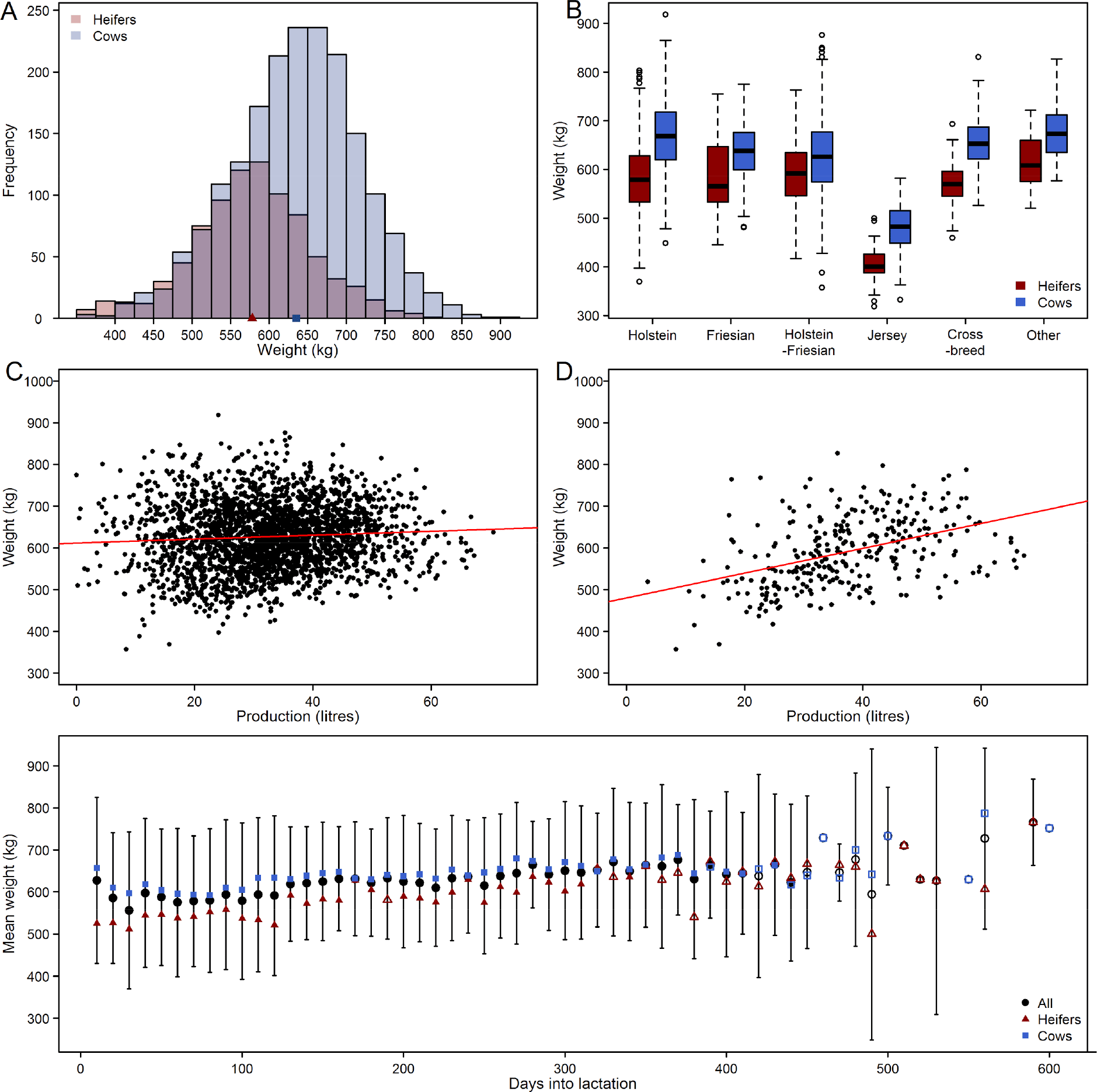
A: Overall distribution of all weights, split into heifers (red) and cows (blue). Mean weights were 578.0 kg for heifers and 635.2 kg for cows, marked with a red triangle and blue square respectively. B: Box plots of the weights of different breeds with heifers (red, left) and cows (blue, right) separated. Heifers were lighter than cows for all breeds (p<0.05, Table 3). C: Scatterplot for weight vs. production in litres for all cattle except Jersey cattle (Pearson’s correlation coefficient = 0.07). Red line indicates line of best fit. D: As for C but for cattle in peak production period only (Pearson’s correlation coefficient = 0.43). E: Trend in weights over the course of lactation for all cattle (black circles, with black lines indicating 95% confidence intervals), heifers (red triangles) and cows (blue squares). Note that confidence intervals are calculated assuming a normal distribution. Points are filled if there are more than 10 cattle at that lactation point, otherwise points are unfilled.

### Effect of breed, lactation number, DIM and production

Some variations in overall mean weight across breeds was seen within the dataset (Table 3, Figure 1B, Figure S1). Of the named breeds, Holstein were the heaviest (636.1 kg) and Jersey the lightest (465.7 kg). Cattle categorised as “Other” were heavier than all breeds (662.8 kg, p<0.01, Table S2).

The proportion of heifers varied between breeds in this dataset. For example, just over 10% of Friesians were heifers, whereas almost 40% of Holsteins were heifers (Figure S2). This is likely to skew the means; indeed, the variation between mean weight of Holstein and Friesian cows was far greater, whereas there was almost no difference between the heifer means for these breeds (Figure 1B, Table S2).

**Table 3.** Summary of the mean weights of breeds represented and relative representation within the UK dairy herd. Note that breed is assigned at farm level. A t-test was used to compare the mean weights of heifers and cows for each breed, and overall

| Breed | N (% heifer) | Overall mean weight, kg (SD) | Heifers mean weight, kg (SD) | Cows mean weight, kg (SD) | p-value |
| --- | --- | --- | --- | --- | --- |
| Holstein | 1,099 (37.9%) | 636.1 (84.1) | 583.9 (73.7) | 668.1 (73.5) | <0.001 |
| Friesian | 130 (13.9%) | 629.3 (65.9) | 586.7 (82.3) | 636.1 (60.6) | 0.024 |
| Holstein-Friesian | 1,099 (26.9%) | 617.4 (72.8) | 590.9 (66.1) | 627.1 (72.8) | <0.001 |
| Jersey | 170 (22.4%) | 465.7 (56.0) | 407.1 (41.9) | 482.6 (47.6) | <0.001 |
| Cross-breeds* | 197 (37.6%) | 623.1 (64.3) | 570.7 (47.0) | 654.6 (51.4) | <0.001 |
| Other ** | 52 (26.9%) | 662.8 (65.8) | 618.4 (56.6) | 679.1 (61.8) | 0.003 |
| Excluding Jersey | 2,577 (31.8%) | 627.3 (77.4) | 585.9 (69.1) | 646.6 (75.0) | <0.001 |
| Overall | 2,747 (31.2%) | 617.3 (85.6) | 578.0 (77.4) | 635.2 (83.2) | <0.001 |

\*cattle recorded as cross-breed
\*\*consisting of different breeds, predominantly dual purpose breeds

There was no correlation between weight and milk production for the 19 Lely farms (production data was unavailable for the Jersey farm) using all cattle (Figure 1C). However, when including only cattle at peak production (days 20-60), cattle with greater production were heavier (Figure 1D).

Mean weight declined for the first thirty days post-calving and was then seen to rise steadily for the remainder of the lactation (Figure 1E). Heifers had a consistently lower weight across lactation than cows.

### Estimated average weight for the UK

Taking the mean weights for different breeds in our dataset (Table 3) and the distribution of these breeds within the UK dairy population (Table S5), we calculated a UK average weight of 619.6 kg.

### Calibration checks

No substantial differences in the mean weight between robots on farms (and hence by breed) were found once proportions of heifers and cows milked by that robot on the day of data collection were accounted for (data not shown to preserve anonymity).

There was little variation in the mean weight for the 6 days of data collected from the single large farm (Figure S3). None of the daily distributions were significantly different from each other (p>0.7) indicating that the calibration of robots was likely to be accurate; significant deviations in weighings from a single robot would affect the distribution and mean weight for that day and would be detected by t-tests (as well as being flagged by the system on farm).

## DISCUSSION

The overall mean weight for all 2,747 dairy cattle was 617.3 kg. Scaling by UK breed proportions gave an estimated average weight for adult UK dairy cattle of 619.6 kg. We therefore suggest a national-level weight of 620 kg to be used for AMU calculations, with farm-level weights to be estimated based on the breed mix on the farm. The most commonly assumed dairy cattle weight in the literature was 600 kg. With our data, we suggest that 600 kg is likely to be an underestimation of mean adult dairy cattle weight in the UK.

There was some variation in weight distribution across all breeds included in this study, ranging from 465.7 kg (Jersey) to 636 kg (Holstein). Jersey cattle were 25.8% lighter than the mean across other breeds. Heifers were on average 9.0% lighter than cows. Cattle categorised as “Other” were heavier than all breeds (662.8 kg, p<0.01, Table S2), however the dataset contained a very low number in this category (n=52, all from 1 farm) and they were predominantly dual-purpose breeds which would be expected to be heavier. The variation between breeds is confounded by differences in the proportions of heifers and cows in each breed. For example, when heifers were removed, the difference in weight between Holstein and Friesian cows widened, though heifers in both breeds had very similar weights. It is possible that Holstein farms may have a tendency to calve heifers at a younger age than Friesian farms. We note that breeds were assigned at the farm level, so it is possible that there was within-farm variation for which we could not account.

Though our sample has a slightly higher proportion of heifers than the NMR data (31.2% compared to 29.1%), as the NMR data could not be split by breed we were unable to use both breed and heifer or cow status accurately in the UK average weight calculation. As heifers weigh less than cows according to our data, this could mean we underestimate the UK average weight.

These data show a decline in mean weight from calving to 30 DIM, and then a steady increase throughout the rest of lactation. These results support trends reported in the literature for both body weight and body condition scores (Dillon et al., 2003, Poncheki et al., 2015). This trend is consistent with the expected period of negative energy balance and the mobilisation of body fat a dairy cow is likely to experience following calving (Eddy, 1992).

Cattle in peak production showed a strong correlation between weight and daily milk production. This may be explained by breed and parity differences: for example, heifers are likely to weigh less and produce less milk than cows, and Holsteins are likely to be higher producers (Holstein UK) and are the heaviest type of named breeds represented in this dataset.

If farms using Lely AMS differed from the average dairy farm, this could create a selection bias. However, the Lely AMS farms used demonstrated a wide variety of management practices and type of cow. The Lely farm animal support advisors were confident that the majority of AMS farms used a ‘standard’ type of dairy cattle, and also stated that many of the farms are flying herds and buying in ‘standard black and white’ cattle as replacements from UK markets. Though there are Jersey farms using Lely AMS which were asked for data, these farms did not record weights. 2.2% of UK cattle are Jersey cattle, which are smaller and lighter than the rest of the UK national herd hence it was important to represent them accurately. This was only possible using an alternative, non-Lely AMS farm, which was weighing cattle using a weigh crush.

All but 3 of the farms used were based in South West England. No data were found to indicate any geographical variation in dairy farming in the UK that would affect our conclusions.

As discussed, Lely robots are calibrated precisely at installation only but are regularly serviced and farmers are advised to regularly clean and tare the weigh floor. This regular cleaning by Lely and the farmer should ensure inaccuracies are minimal. During data collection for this project, farmers were asked to calibrate the scales. The normal distribution of the data indicates that there were no major inaccuracies unless identical inaccuracies were occurring on every farm in the dataset, which seems unlikely. Indeed, data obtained over a week from a single farm showed no significant difference in mean weight between days.

**Figure 2:**
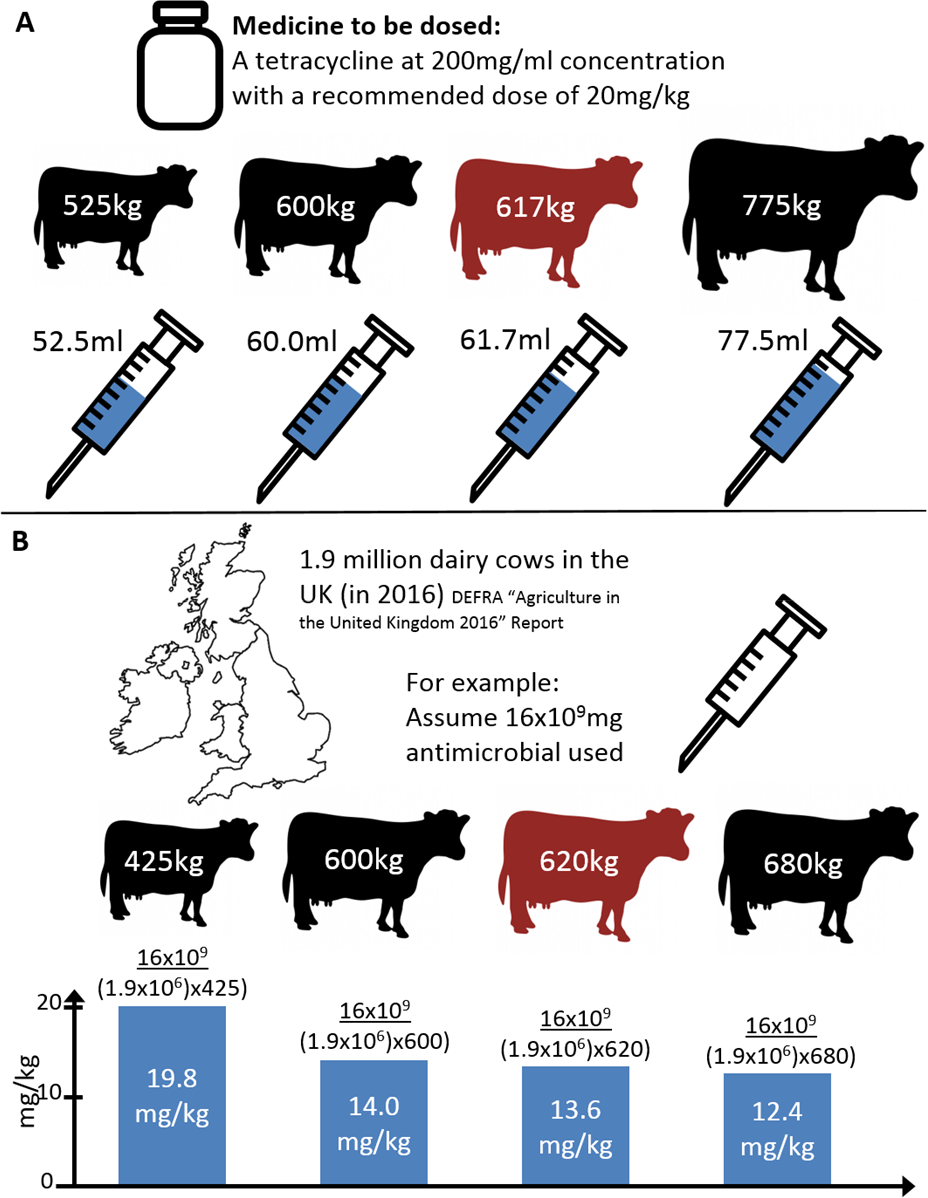
Illustration of the effect different assumed cattle weights can have on the medicine dose for Holstein-Friesians (panel A) and effect on the resulting mg/kg metric when measuring antimicrobial use in dairy cattle (panel B). In panel A, 525 and 775 kg weights were the lowest and highest estimates from practising veterinarians asked to estimate the average weight of a UK Holstein-Friesian milking cow, 600 kg was the most common adult dairy cattle weight reported (Table 1), and 617 kg was the mean weight of Holstein-Friesians estimated in this work. In panel B, note that a usage of 16×10^9^ mg of antimicrobial in the UK is intended as an example only. 425 kg was the lowest dairy cattle weight reported in the literature (as the “estimated weight at time of treatment” (European Surveillance of Veterinary Antimicrobial Consumption)), 600 kg was the most common weight reported (Table 1), 620 kg was the UK mean weight estimated in this work and 680 kg was the most extreme weight reported in the literature (Table 1, note this weight was from the USA (Pol and Ruegg, 2007)).

This study is the first to estimate a mean weight of UK dairy cattle based on data. Weights from 2,747 cattle from the 4 main named breeds, as well as cross<u>--</u>breeds and less common breeds were considered. These data provide valuable evidence to support 620 kg as an appropriate average weight of UK adult dairy cattle. The impact of having an evidence-based figure for the average weight, as well as variation by breed, production level and days in milk will be marked. For example, for dosing, visual weight estimation of individual cattle will be easier and more accurate if an actual average is known in the first instance (Figure 2A). Also, for farm-level and national-level antimicrobial use reporting, our recommended UK weight of 620 kg will be invaluable, as using too high or too low a weight can significantly impact calculations of antimicrobial use (Figure 2B).

## ACKNOWLEDGEMENTS

The authors would like to thank Wendy Ward-van Winden, Farm Management Support Advisor at Lely Center, Holsworthy for her help in obtaining data and providing additional information relating to Automatic Milking Systems. Thanks also to Jon Eldridge, Farm Management Support at Lely Center, Yeovil and Bas van Senten, Farm Management support manager, UK and Ireland for their help in obtaining data. Thanks also go to all farmers who contributed data to the study, in particular Matthew Davey who provided additional data and answered many questions. We thank Fraser Broadfoot of the Veterinary Medicines Directorate for providing data on cattle breeds in the UK. Thanks also to the Farm Animal Group at Bristol Veterinary School for their comments on early drafts of this work, and to Emma Wright for assisting with data collection. Thanks to Kiera Schubert of Torch Farm Vets for comments on the final draft. Finally, we thank Seamus Gilheany, NMR Software Development at National Milk Records for providing data on some basic statistics of a sample of the UK national dairy herd.

